# Chromosome-Scale Genome Assembly Provides Insights into Speciation of Allotetraploid and Massive Biomass Accumulation of Elephant Grass (*Pennisetum purpureum* Schum.)

**DOI:** 10.1101/2020.02.28.970749

**Authors:** Shengkui Zhang, Zhiqiang Xia, Wenqing Zhang, Can Li, Xiaohan Wang, Xianqin Lu, Xianyan Zhao, Haizhen Ma, Xincheng Zhou, Weixiong Zhang, Tingting Zhu, Pandao Liu, Guodao Liu, Hubiao Yang, Jacobo Arango, Michael Peters, Wenquan Wang, Tao Xia

## Abstract

Elephant grass (*Pennisetum purpureum* Schum., A’A’BB, 2n=4x=28), which is characterized as robust growth and high biomass, and widely distributed in tropical and subtropical areas globally, is an important forage, biofuels and industrial plant. We sequenced its allopolyploid genome and assembled 2.07 Gb (96.88%) into A’ and B sub-genomes of 14 chromosomes with scaffold N50 of 8.47 Mb. A total of 38,453 and 36,981 genes were annotated in A’ and B sub-genomes, respectively. A phylogenetic analysis with species in *Pennisetum* identified that the speciation of the allotetraploid occurred approximately 15 MYA after the divergence between *S.italica* and *P. glaucum*. Double whole-genome duplication (WGD) and polyploidization events resulted in large scale gene expansion, especially in the key steps of growth and biomass accumulation. Integrated transcriptome profiling revealed the functional differentiation between sub-genomes; A’ sub-genome contributed more to plant growth, development and photosynthesis whereas B sub-genome primarily offered functions of effective transportation and resistance to stimulation. The results uncovered enhanced cellulose and lignin biosynthesis pathways with 645 and 666 genes expanded in A’ and B sub-genomes, respectively. Our findings provided deep insights into the speciation and genetic basis of fast growth and high biomass accumulation in the species. The genetic, genomic, and transcriptomic resources generated in this study will pave the way for further domestication and selection of these economical species and making them more adaptive to industrial utilization.

## Introduction

Since the Cenozoic era, the adaptation of the biosphere to high temperature, drought and the increase of CO2 concentration have led to the expansion of C4 plants, and as the result greatly increased the bioaccumulation, which is critical for forage livestock and biomass energy utilization^1,2^. Example species of these C4 plants include elephant grass (*P. purpureum* Schum.) and pearl millet *(P. glaucum*) in the *Pennisetum* genus. This genus has about 140 members that are broadly distributed in different environments around the world^3,4^. Members of the *Pennisetum* genus are generally characterized with fast growth, high temperature and drought tolerance, and high biomass accumulation.

Elephant grass naturally distributed or cultivated in semi-arid tropical and sub-tropical regions of Africa, Asia, and America^5,6^. It is an economically important tropical forage crop, which provides a considerable amount of valuable grazing grasses for ruminant cattle and sheep in the regions. Elephant grass is also an excellent material for producing biofuel, biochar, alcohol, methane, and paper^7–10^. It has been characterized as fast growth, tolerance to abiotic and biotic stresses and huge biomass yielding potential ^11^. Under desirable growth conditions, elephant grass could grow to a height of 2-6 meters with huge biomass productivity of about 45 t/ha and can be harvested 3-4 times yearly^12,13^.

The elephant grass species (A’A’BB, 2n=4x=28) was originated in East Africa by natural hybridization between two diploid progenitors, the pearl millet (*P. glaucum*, AA, 2n=14) and an unknown species, according to fossil and cytogenetic clues^14^. Pearl millet has been domesticated as an edible grain that is cropped specifically in arid regions in Africa and India where the other grains failed to reproduce seeds because of too little rainfall. Elephant grass has an artificial triploid offspring king grass (*P. purpureum* × *P. glaucum*) (AA’B, 2n=21), which is 20% higher yield widely cultivated in tropical regions around the world.

Elephant grass is an allotetraploid crop with a complex genome. The species is primarily cross-pollinated because of its androgynous flowering behavior, resulting in high heterozygosity and large genetic diversity which can be utilized in breeding programs. The genetic research of elephant grass has been so far mainly focusing on evaluating genetic diversity through constructing molecular markers and fingerprints and determining genetic relationship^15–17^. Recently, a ~1.79 Gb draft whole-genome sequence of pearl millet was acquired by the whole-genome shotgun (WGS) and bacterial artificial chromosome (BAC) sequencing^18^. The relationship between elephant grass (A’A’BB) and pearl millet (AA) was clarified by genomic *in situ* hybridization (GISH)^19^. A high degree of homology between genomes A and A’ was confirmed, whereas genome B presented lower percentage marks that were observed only in the centromeric and pericentric regions of all chromosomes of A’ genome.

Despite the amount of effort into the research of the *Pennisetum* genus, its genomic and genetic resources, particularly that for elephant grass and its relatives, have been scarce so far. It is now essential and urgent to acquire a high-quality genome assembly of elephant grass for understanding the evolution of species in the genus and deciphering the genetic underpinning of many physical and economical traits of this important forage crop, including the fast growth, stress tolerance, and biomass accumulation. In this study, the elephant grass cultivar CIAT6263 was chosen for genome sequencing using third-generation sequencing technologies. Single-molecule long-read sequencing technologies including Pacific Biosciences (PacBio) and Oxford Nanopore Technologies (ONT) are able to reveal complex genome structures^20,21^. These techniques have been adopted to construct high-quality assemblies of large genomes, such as human^22^, *Arabidopsis thaliana*^23^, *Solanum pennellii*^24^ and sorghum^25^. In addition, long-range scaffolding technologies, such as Bionano Genomics optical maps and high-throughput chromatin conformation capture (Hi-C) proximity ligation^26^, in combination with long-read ONT sequences can generate chromosome-scale scaffolds, which have been applied to sequence the genomes of sorghum^25^, *Brassica rapa*^27^, and *Spirodela polyrhiza*^28^.

Using an integrative strategy that combined Nanopore, Bionano optical physical map, and Hi-C proximity ligation, we constructed a high-quality, chromosome-level allopolyploid genome of elephant grass cultivar CIAT6263 and annotated the genes into the A’ and B sub-genomes. Furthermore, exploiting the rich genomic resources we constructed the phylogenetic tree of the species in the *Poaceae* family, which helped reveal the double whole-genome duplication (WGD) events that resulted in polyploidization and the expansion of gene families in elephant grass. Such large scale genomic reorganization could set the genetic bases for the key development supporting fast growth, drought tolerance, and high biomass. In particular, the genetic mechanism of cellulose and lignin synthesis in elephant grass was closely examined at the level of whole-genome and gene family. Together with transcriptome profiling, we revealed the functional differentiation between A’ and B sub-genomes. This work provides a solid foundation for understanding the genetics of the extraordinary traits and evolutionary trajectory in species of *Penniesum*. Furthermore, the precision assembly and annotation of an allotetraploid genome of more than 2.0 Gb contributes a new example for the sequencing of species with super large polyploid genomes.

## Results

### De novo Assembly of Allotetraploid Elephant Grass Genome

The genome size of elephant grass accession CIAT6263 (Supplementary Fig. 1a) was estimated to be 2.0-2.13 Gb by a k-mer analysis and flow cytometry (Supplementary Fig. 2 a-b and Supplementary Table 2). A total of 8.85 million Nanopore clean reads (~186.35 Gb data, ~93× coverage) and 1.70 million clean ultra-long reads (63.99 Gb data, 32× coverage) were generated using Nanopore Technologies system (Supplementary Table 1). Initial assembly yielded 2.07 Gb of assembled sequence with contig N50 2.9Mb (Table 1).

**Table 1.**
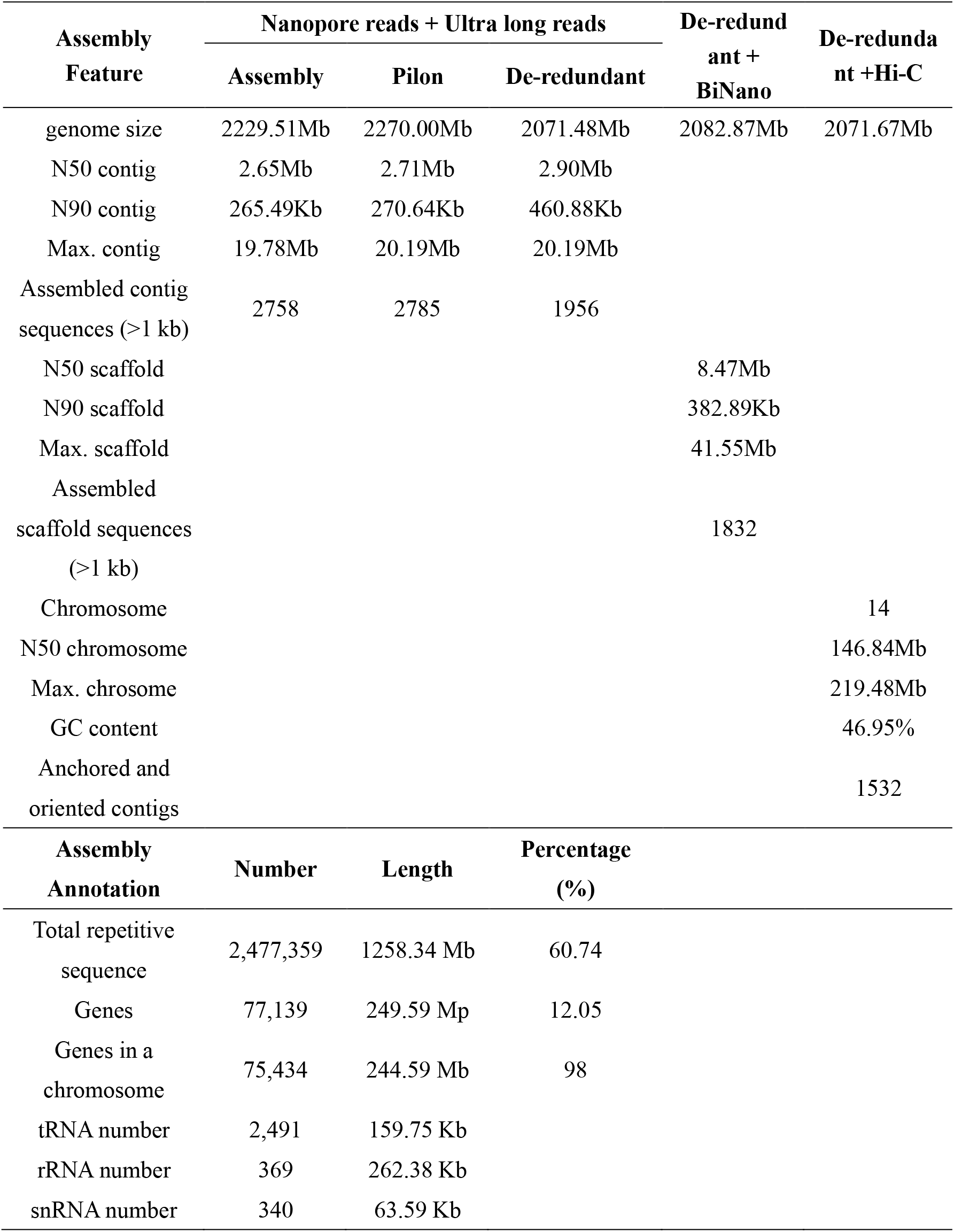
Overview of Genome Assembly of Elephant Grass and Gene Predicition.

| Assembly<br>Feature | Nanopore reads + Ultra long reads |  |  | De-redund<br>ant +<br>BiNano | De-redunda<br>nt +Hi-C |
| --- | --- | --- | --- | --- | --- |
|  | Assembly | Pilon | De-redundant |  |  |
| genome size | 2229.51Mb | 2270.00Mb | 2071.48Mb | 2082.87Mb | 2071.67Mb |
| N50 contig | 2.65Mb | 2.71Mb | 2.90Mb |  |  |
| N90 contig | 265.49Kb | 270.64Kb | 460.88Kb |  |  |
| Max. contig | 19.78Mb | 20.19Mb | 20.19Mb |  |  |
| Assembled contig<br>sequences (>1 kb) | 2758 | 2785 | 1956 |  |  |
| N50 scaffold |  |  |  | 8.47Mb |  |
| N90 scaffold |  |  |  | 382.89Kb |  |
| Max. scaffold |  |  |  | 41.55Mb |  |
| Assembled<br>scaffold sequences<br>(>1 kb) |  |  |  | 1832 |  |
| Chromosome |  |  |  |  | 14 |
| N50 chromosome |  |  |  |  | 146.84Mb |
| Max. chrosome |  |  |  |  | 219.48Mb |
| GC content |  |  |  |  | 46.95% |
| Anchored and<br>oriented contigs |  |  |  |  | 1532 |
| Assembly<br>Annotation | Number | Length | Percentage<br>(%) |  |  |
| Total repetitive<br>sequence | 2,477,359 | 1258.34 Mb | 60.74 |  |  |
| Genes | 77,139 | 249.59 Mp | 12.05 |  |  |
| Genes in a<br>chromosome | 75,434 | 244.59 Mb | 98 |  |  |
| tRNA number | 2,491 | 159.75 Kb |  |  |  |
| rRNA number | 369 | 262.38 Kb |  |  |  |
| snRNA number | 340 | 63.59 Kb |  |  |  |

The assembly was optimized and modified by 329.53 Gb (150X) BioNano long-reads clean data, resulted in a 2.07 Gb genome with a scaffold with N50 of 8.4 Mb (Table 1 and Supplementary Table 1,3). The contigs and scaffolds were further scaffolded into 14 chromosomes anchored 2.003 Gb (96.88%) of the genome by the Hi-C technology (Fig. 1, Table 1, Supplementary Fig. 3, Supplementary Table 1, 3).

**Figure 1.**
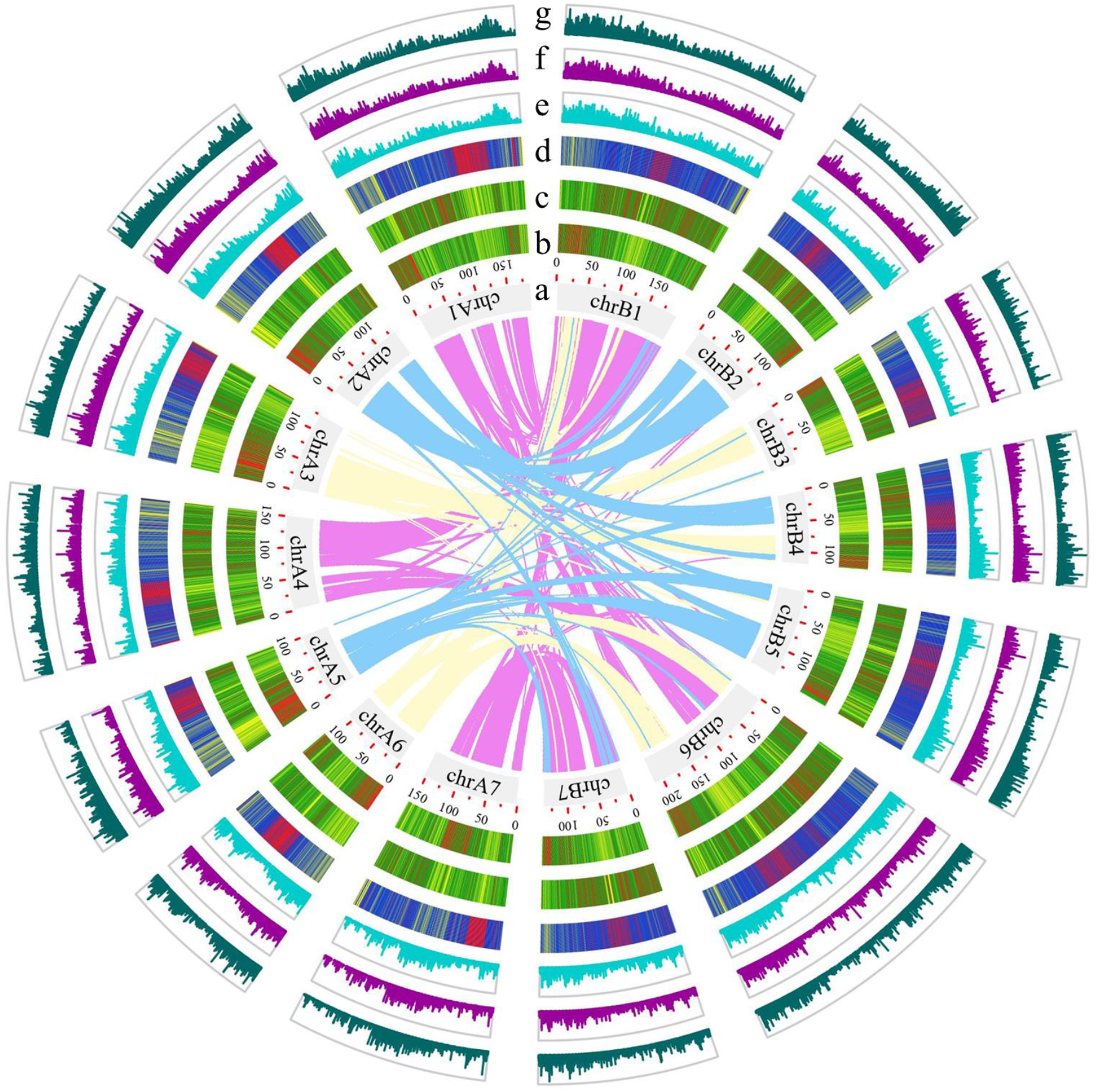
Overview of the Allotetroploid Elephant Grass Genome. The genome was assembled into A’ and B sub-genomes each with 7 chromosomes. The Circos plot of the multidimensional topography from outermost to innermost, a-f, shown that: (a) the 14 chromosomal pseudomolecules, units on the circumference are megabase values of pseudomolecules, (b) gene density, (c) repeat density, (d) 5mC DNA methylation levels, and gene expression levels in(e) root, (f) stem and (g) leaf(b, c, d) are shown in 1 Mb windows sliding 200 kb, (e, f, g) are shown in 50 kb windows sliding 5 kb. Central colored lines represent syntenic links between A’ and B sub-genomes.

By comparing with the pearl millet (*P. glaucum*, AA, 2n=14) genome sequences^18^, we successfully sorted the genome of elephant grass into A’ and B sub-genomes with 14 chromosomes (Fig.1a and Supplementary Fig. 4). The assembled genome was more than 97.8% complete (Supplementary Table 4) when evaluated using BUSCO^29^. The alignment of the assembled transcripts from RNA sequencing (RNA-seq) with the genome revealed an approximately 90% sequence identity (Supplementary Table 7).

### Genome Annotation

The genome annotation was performed using the AUGUSTUS pipeline^30^ incorporating *ab initio* predictions and transcriptome data, resulting in 77,139 protein-coding genes (Table 1). Of the 77,139 genes, 75,434 (98%) were assigned to chromosomal locations, including 38,453 in A’ sub-genome and 36,981 in B sub-genome (Table 1 and Supplementary Table 5). These genes were unevenly distributed along the chromosomes with a distinct preference for the ends (Fig. 1b). We also annotated 369 ribosomal RNAs (rRNAs), 2491 transfer RNAs (tRNAs) and 340 small nuclear RNAs (snRNAs) (Table 1). Meanwhile, 1.26 Gb of repeat elements were identified in the 2.07-Gb genome assembly, showing that 60.74% of the assembled genome was repetitive (Table 1 and Supplementary Table 5). These repeat elements were unevenly distributed along the chromosomes with a distinct preference to the centromeres (Fig. 1c). In common with the pattern in many other plant genomes, long-terminal repeat (LTR) retrotransposons were the most abundant class of the repetitive DNA. It accounts for more than 42% of the nuclear genome of elephant grass, among which Gypsy repeats being the most abundant, followed by Copia (Supplementary Table 6). Among these repeat elements, the proportion on B sub-genome (65.71%) was higher than that on A’ sub-genome (60.59%), and the same was true for the types of LTR and SINE (Supplementary Table 5, 6). The differences between the distributions of genes and repeats across the genome may account for the different functions of the A’ and B sub-genomes.

Additionally, 2,910 transcription factors (TF) in 89 families, 2,437 protein kinases (PK) in 118 families were identified separately (Supplementary Table 8, 9). Elephant grass possessed more TFs than *P. glaucum*, especially the TF families of *B3*, *bZIP*, *C3H*, *FAR1* and *NAC* (Supplementary Table 8), which may play a pivotal role in growth and development. It also had more PKs than *P. glaucum* (1,199). For example, there were 1,660 RLK-Pelle like protein kinases, especially 585 members in the DLSV subfamily which were substantially more than that in pearl millet and other 5 species (Supplementary Table 9), involved in disease immunity and regulation of growth and development^31^.

### Phylogenetics of the Allotetraploid Elephant Grass and *Pennisetum* Species

Except for elephant grass, eight other economically important species in *Gramineae* have been sequenced. *P. glaucum*, *S. italica*, *P. miliaceum*, *Z. mays*, and *S. biolor* belong to *Panicoideae*, and *O. sativa*, *B distachyon*, and *A. tauschii* ascribed into *Poaceae*. A phylogenetic tree was constructed using a set of unique gene families among above 8 relative species and some other 4 outer species (Fig.2a and Supplementary Table 10). The species in *Pennisetum* diverged approximately 22 million years ago (MYA) from *Setaria* which was referenced as *S. italic* and the pearl millet (*P. glaucum*) diverged about 20 MYA. The allotetraploid elephant grass with A’A’BB genome was originated about 15 MYA. It was perhaps caused by the grazing pressure while herbivores outbreak in the Miocene era of Africa^18,32^.

**Figure 2.**
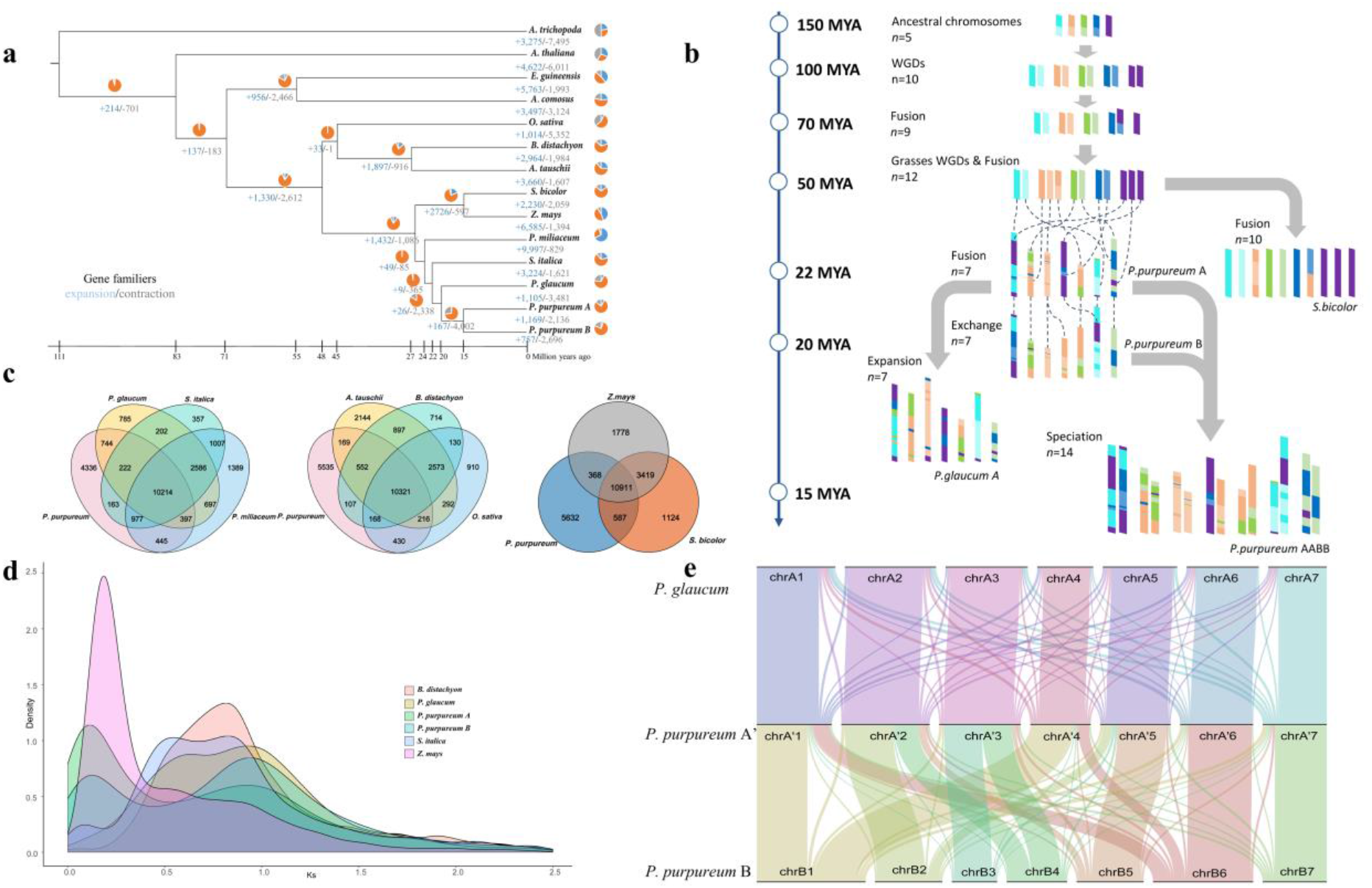
Evolution of Elephant Grass and Relative Species in *Pennisetum*. a. Phylogenetic tree of 12 species and A’/B sub-genome of elephant grass. Gene family expansion and contraction compared with the most recent common ancestor. Gene family expansions are indicated in blue. Gene family contractions are indicated in gray. Inferred divergence times (MYA, million years ago) are denoted at each node. Venn diagram shows the ratio of gene family expansions and contractions. b. Modern chromosome derivation in A’/B sub-genome of elephant grass, *P. glaucum* from ancestral chromosomes. c. Shared and unique gene families in different species of *Poaceae*. d. Distribution of Ks values of the best reciprocal BLASTP hits in the genomes of *P. purpurum* A’, *P. purpurum* B, *P. glaucum*, *S. italica*, *B. distachyon* and *Z. mays*. e. Syntenic blocks between A’/B sub-genome of elephant grass and *P. glaucum.*

Furthermore, we constructed the evolutionary trajectory of the species in *Panicoideae,* especially those in *Pennisetum,* according to the relationship of their orthologous genes (Fig. 2b and Supplementary Table 11). The reconstruction of the ancestral chromosome of grasses has revealed that the ancestor had 12 chromosomes (referenced as rice) after one WGD and two nest chromosome fission events^33^. Using rice as a reference to investigate the chromosome evolution of elephant grass, all chromosomes of A’ sub-genome except Chr2 were colinear with rice chromosomes (Supplementary Fig. 5c). Most of the chromosomes(Chr3, 4, 5, 6 and Chr7) of B sub-genome were collinear with rice chromosomes(Supplementary Fig. 5d). All chromosomes of *P. glaucum* were consistent with rice (Supplementary Fig. 5b).

The comparison of A’ and B genomes with the genomes of foxtail millet (*S.italica*) and *B. distachyon* independently showed that all seven chromosomes in A’ or B sub-genome of elephant grass corresponded strongly to the nine chromosomes of foxtail millet and the five chromosomes of *B. distachyon* (Supplementary Fig.6). The highly conserved collinearity supported that there existed a close evolutionary relationship among these species. This suggested that fusions and divergence might have occurred among the chromosomes of the ancestors of these species in the processes of evolution and adaptation to the environment.

It is found that the multiple fusion of chromosomes took place 22 MYA, thus forming the species of *Pennisetum* (n=7) (Fig. 2b). Selection and domestication drove the speciation of pearl millet (*P. glaucum*, AA) and elephant grass (Fig. 2b). The evolution of A’ and B sub-genomes in the genus was also clarified by the collinear analysis (Fig. 2e). The A’ sub-genome of elephant grass was highly paralogous with A genome of pearl millet, except for slight gene flow among the 7 chromosomes each other. ChrB1 integrated with ChrA1, A4 and some parts of A6 and A7, ChrB2 corresponded to ChrA2 and A5, ChrB4 had synteny with ChrA2 and ChrA3, and ChrB6 was related to ChrA1, A4 and A6. A’ and B sub-genomes of elephant grass and A genome of pearl millet shared common ancestor chromosomes, and the differences were caused by fusion and rearrangement of chromosomes.

### Genome Duplication

We investigated probable genome duplication events of elephant grass using synonymous substitutions (Ks) density plots of orthologous and paralogous gene pairs which were selected from A’ and B sub-genomes of elephant grass and relative genomes. It was evident that there were two peaks for both A’ and B sub-genomes (Fig. 2d), the peak Ks 0.9 shared with grass suggested an ancient WGD event which happened around 50 MYA (Fig. 2d), the similar peak Ks 0.2 indicated that the modern WGD event occurred in A’ and B sub-genomes around 15 MYA. It was consistent with the results of the phylogenetic tree. Adding the speciation of allotetraploid with the integration of A’ and B genome formed in that time (Fig. 2a,b, Supplementary Fig.7), the elephant grass genome probably went through polyploidy three times. Therefore, ancient WGDs in elephant grass preceded before it diverged from foxtail millet and *B. distachyon* (Supplementary Fig.7).

### Expansion of Gene Families and Gene Copy Numbers

The enhanced and shared gene families among A’ and B sub-genomes and 12 other species (Supplementary Table 10) were identified using OthoMCL^34^. Elephant grass shared 10,214 gene families with close relatives *P. glaucum*, *S. italic*, and *P. millaceum*, 10,911 gene families with *Z. mays* and *S. bicolo*r, and 10,321 gene families with *B. distachyon*, *O. sativa*, and *A. tauschii* (Fig. 2c). Among all these species, 6,598 gene families were shared, whereas 2,733 and 2,266 gene families were specific to A’ and B sub-genome of elephant grass, respectively (Supplementary Table 12). It indicated that more unique gene families were enhanced in elephant grass than in the other seven species in the grass family (Fic. 2c). Gene Ontology (GO) annotation showed that these genes were enriched in transporter activity, catalytic activity, cellular process, and metabolic process in A’ and B sub-genomes (Supplementary Fig. 8a-b). In reference to A genome of pearl millet, 4,568 new gene families were specific to A’ sub-genome and 5,921 gene families were unique in the elephant grass genome, showing a huge expansion of gene family. These genes also had similar functions described above (Fig. 2c, Supplementary Fig. 10, 12 a-b).

Expansion and contraction of gene families are an important feature of the selective evolution of species^35^. In comparison with the evolutionary nodes of genome A of pearl millet, it was found that 1,169 gene families expanded substantially and 1,282 contracted in A’ sub-genome whereas 757 gene families expanded and 1,699 contracted in B sub-genome of elephant grass (Fig. 2a). Significant expansion occurred in some gene families in A’ and B sub-genomes, with gene copy numbers varying from 2 to 74. All the highly expended gene families in A’ genome were ascribed into chromatin regulatory components, transcription factors (TFs), kinases, and transporters, except for a small number of unknown functional proteins. The highly expanded gene families in B sub-genome were slightly enriched in transport relative proteins, TFs and Kinases. GO annotation found that the gene families contracted in A’ sub-genome were enriched in binding and transcription regulator activity process, whereas in B sub-genome the same gene families were enriched in the cell, cell part, organelle, macromolecular complex, transcription regulator activity and binding (Supplementary Fig. 9a-d). The selective expansion or contraction of these gene families which were preserved in evolution suggested the important environmental adaptation of the species.

Comparing with pearl millet (A genome), 2,532 and 8,873 gene families expanded in A’ sub-genome and elephant grass genome, respectively. The average copy number of the expanded gene families in A’ sub-genome is 4.23 (range of 2~190) which was greater than 1.73 (range of 1~49) of that in pearl millet, whereas 4.15 (range of 2~192) in elephant grass was also significantly greater than 1.64 (range of 1~64) in pearl millet (Supplementary Fig. 11). Among the expended gene families, 3,919 with 2 copies were probably resulted from the heteropolyploidy of the single-copy genes of elephant grass. Meanwhile, 1,835 and 781 gene families contracted in A’ sub-genome and elephant grass, respectively. More gene families expanded than contracted over the evolution of elephant grass. GO annotation showed that those significantly expanded gene families in A’ sub-gnome or elephant grass genome were mainly enriched in cell and cell part, cellular process, metabolic process, catalytic activity and binding, biological regulation and response to stimulus (Supplementary Fig. 12 c-d).

### Differentiation of Biological Functions in A’ and B Sub-genome

We annotated 38,453 and 36,891 genes in A’ and B sub-genomes, respectively (Supplementary Table 5, 12). Among them, 12,499 were orthologous gene families, 27,33 were specific to A’ sub-genome, and 2,266 specific to B sub-genome, including 11,257 gene families in A’ and B sub-genomes exactly matched (Supplementary Fig. 10 and Supplementary Table 12, 13). All the homologous genes were assigned into 19 biological processes (mainly cellular and metabolic processes), 6 cellular components (mainly organelle, cell, and cell part) and 5 molecular functions (mainly catalytic activity and binding). There was no evident biased functional divergence in A’ and B sub-genomes (Supplementary Fig. 13a). The gene families specific to B sub-genome were enriched mainly in transport activity, especially in substrate-specific transporter activity, transmembrane transporter activity, and sugar symporter activity. In contrast, no significant enrichment was detected in the gene families specific to A’ sub-genome (Supplementary Fig. 13a-c).

We collected 19 samples of vegetative organs of leave and different notes of stem and root at three developing stages (Supplementary Fig. 1b-d and Supplementary Table 7) and used them for transcriptome profiling and annotation using deep sequencing. The different expression profiles of genes in A’ and B sub-genomes in various organs strongly supported the diversification of biological functions in the two sub-genomes (Fig. 1e-g, Supplementary Fig. 14 and Supplementary Table 13,14). In the homologous gene families, genes from A’ sub-genome mainly ascribed into chloroplast and plastid related which were involved in energy conversion and storage (in leaf), cell part and intracellular, indicating the important role of A’ sub-genome in promoting stem development (in stem) and plasma membrane (in root) (Fig. 3I a-f). However, genes from B sub-genome were mainly related to various stimulus resistance, maintaining the normal development in adverse environment (in leaf), chemical stimulus-response and transmembrane transporter activity (Fig. 3III h-m). The functions of differentially expressed genes (DEGs) in two sub-genomes were conspicuously different but complementary to each other to promote growth and development adaptable to diverse environments. It was in accordance with the previous results that the unequal expression of homologous genes in allopolyploids can be an important feature and consequence of polyploidization^36,37^.

**Figure 3.**
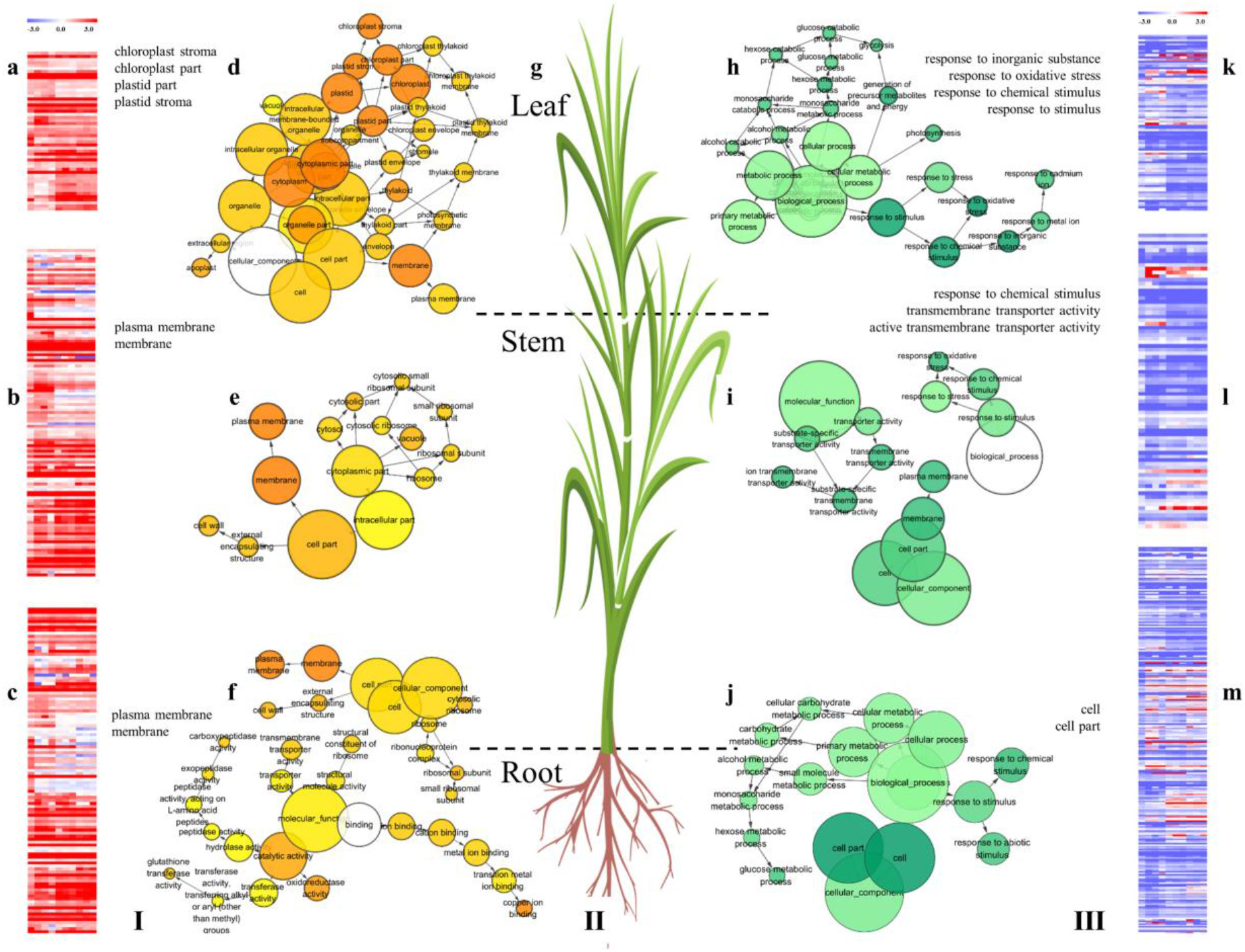
Homologous Genes of A’ and B sub-genome in Elephant Grass. The heat map and GO annotation of homologous in A’ sub genome (Ⅰ, a-f) and B (III, h-m) in leaf, stem and root; a,b,c,k,l,m. The heat map shows the log2-based A’/B TPM changes fold value in leaf (a, k), stem (b. l), root (c, m), red indicated homologous in A’ genome have high expression, blue indicated homologous in B genome have high expression. The different tissues from left to right were T1_S1, T1_S2, T1_S3, T2_S1, T2_S2, T2_S3, T2_S4, T3_S1, T3_S2, T3_S3, T3_S4, T3_S5, T3_R, T3_L1_H, T3_L1_M, T3_L1_T, T3_L2_H, T3_L2_M, T3_L2_T. d,e,f,h,i,j. Significantly enriched biological process Gene Ontology (GO) categories (yellow, A’ subgenome; green, B subgenome) in leaf (d, h), stem (e, i) and root (f, j). Color intensity reflects significance of enrichment, with darker colors corresponding to lower P values. Circle radii depict the size of aggregated GO terms. II(g). The pattern of elephant grass, divided into leaf, stem and root.

### Enhanced Cellulose Synthesis Metabolism in Elephant Grass

Well known for its extremely high biomass yield, elephant grass consisted of cellulose (36%), hemicellulose (23%), lignin (13%) and other non-structural components. The gene families especially the plasma membrane-localized cellulose synthase (*CesA*), *SUS*, *CINV*, and *HXK* in the cellulose synthesis pathway were highly expanded (Fig. 4a). We detected 56 cellulose synthase (*CesA*) and cellulose synthase-like (*Csl*) genes in elephant grass, representing higher copy numbers than *A. thaliana* (40) and *O sativa* (43)^38^. Tissue expression analysis showed that 30 *CesA* genes had high expression levels, while 11 genes had low expression levels, and the expression levels in stems were higher than that in roots and leaves. Meanwhile, there were 17 *SUS*s, 6 *UDGP*s, *CINV*s and *HXK*s which were expanded and more highly expressed in leaves and stem than in roots (Fig. 4a and Supplementary Table 15). The results of the Q-PCR analysis of some genes were consistent with the result from transcriptome profiling (Supplementary Fig. 15a). The origin of the expanded gene members in each family was investigated. They were normally distributed in corresponding chromosomes of A’ and B sub-genomes, and two and three tandem duplicates appeared in A’ and B sub-genomes (one of which is three tandem genes), respectively (Fig. 4c). These results indicated that A’ and B sub-genomes contributed equally to highly efficient cellulose biosynthesis toward massive biomass accumulation.

**Figure 4.**
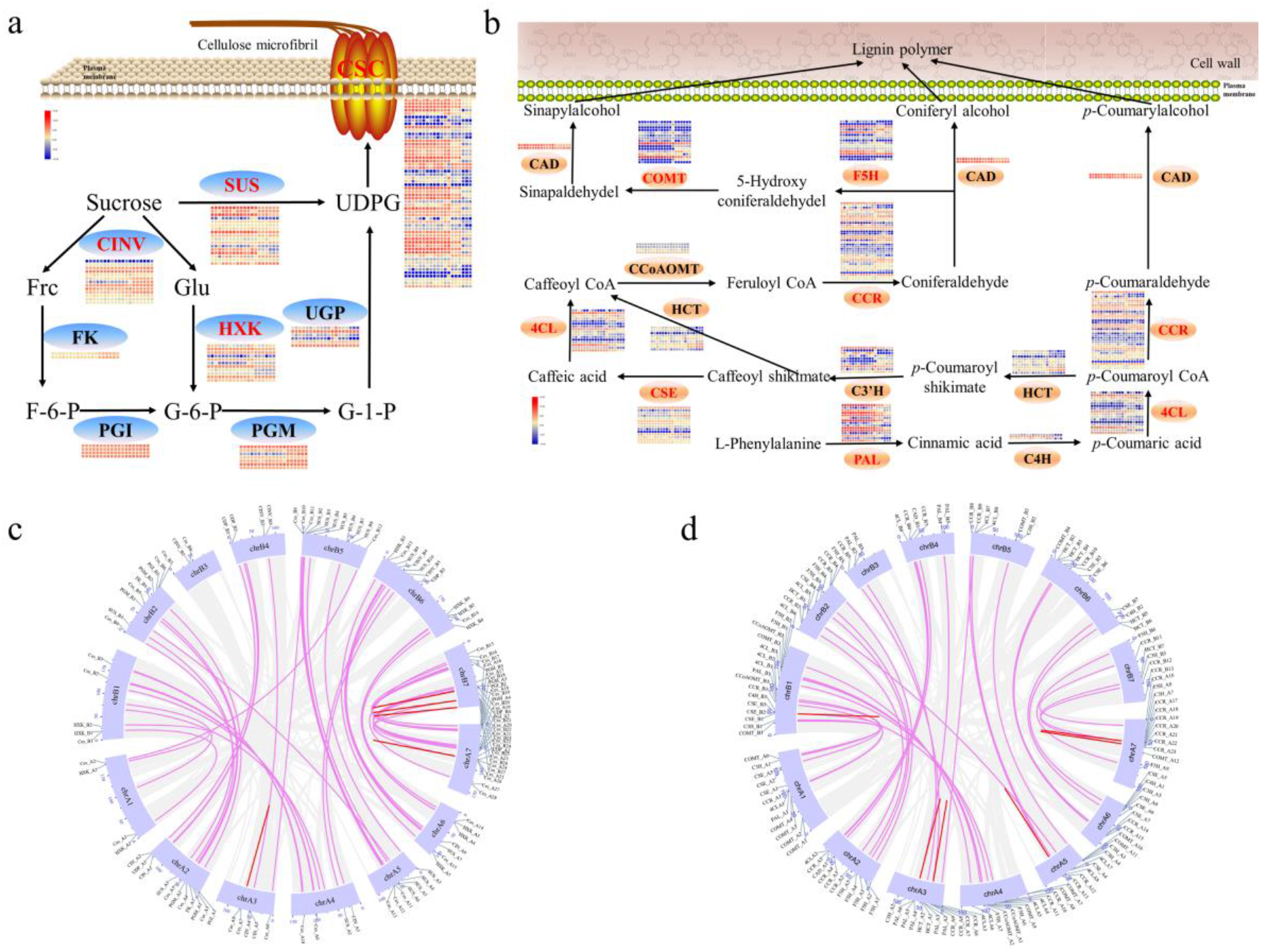
Gene Expansion and Expression Heatmap of Cellulose and Lignin Synthesis in Elephant Grass. a,b. Overview of the cellulose biosynthesis (a) and lignin biosynthesis (b) pathway shown the gene number expansion in each step and their expression profiling in organ of vegetation.^69^. Red is high and blue is low. The different tissues from left to right were same as figure 3. The heatmap was drawn using log2-based TPM-changed fold values. The differential expression thresholds of significant up and down regulation were 15.0 and –10.0, respectively. The gene name in red indicates expansion: *CSC*, cellulose synthase complex; *CINV*, cytosolic invertase; *FK*, fructokinase; *HXK*, hexokinase; *PGI*, phosphoglucoisomerase; *PGM*, phosphoglucomutase; *SUS*, sucrose synthase; *UGP*, UDP-glucose pyrophosphorylase; *4CL*, 4-coumarate CoA ligase; *C30H*, p-coumaroyl shikimate 30-hydroxylase; *C4H*, cinnamate 4-hydroxylase; *CAD*, cinnamyl alcohol dehydrogenase; *CCoAOMT*, caffeoyl CoA O-methyltransferase; *CCR*, cinnamoyl CoA reductase; *COMT*, caffeic acid O-methyltransferase; *CSE*, caffeoyl shikimate esterase; *F5H*, ferulate 5-hydroxylase; *HCT*, hydroxycinnamoyl CoA shikimate hydroxycinnamoyl transferase; *PAL*, phenylalanine ammonia lyase. c,d. Synteny analyses of gene in Cellulose biosynthesis (c) and Lignin biosynthesis (e) between A’ - and B-genomes. Purple lines indicate homologous gene pairs, red lines indicate tandem duplication, gray strips indicate aligned syntenic blocks. The gene in the outer ring represents location of genes.

In addition, WGCNA analysis^39^ was performed on the data from the 19 samples and the result was divided into 9 modules (Supplementary Fig. 16). As the modular network heatmap on the genes related to lignin and cellulose synthesis shows (Supplementary Fig. 17), 644 genes were involved in cellulose synthesis, 28 of which were transcription factors such as *MYB*, *bZIP,* and *MADS*. Among the 644 genes, 300 formed 9 gene modules and the expression trend was consistent in different samples (Supplementary Fig. 18a). The gene expression networks for the leaf and root were different and there was no significant difference in gene expression for the genes in A’ and B sub-genomes. (Supplementary Fig. 17a-b).

### Strengthened Lignin Synthesis in Elephant Grass

We annotated all gene families in the lignin biosynthesis pathway and found gene expansion in key gene families (Fig. 4b). In total, elephant grass possessed 135 lignin biosynthesis-related genes (Supplementary Table 16), and in comparison, *O. sativa* had 109 and *P. heterocycle* had 87. The higher gene number of elephant grass may be due to the doubling of tetraploid speciation. Gene families such as *PAL*, *4CL*, *F5H*, *COMT*, *CSE,* and *CCR* were expanded. The members of genes in these gene families ranged from 2 to 36, and the genes were located mainly on chromosomes 1, 2, 4, 6 and 7 of A’ and B sub-genomes. More lignin synthesis related genes existed on A’ sub-genome than on B sub-genome. We also found that 38 homologous gene pairs and 5 tandem repeat regions in A’ and B sub-genomes (Fig. 4d). In addition, 664 genes involved in lignin synthesis were identified by WGCNA analysis, 23 of which were transcription factors such as *MYB*. Among them, 271 genes constituted 6 gene modules, and the expression trend of different samples was consistent (Supplementary Fig. 17 b-d and Supplementary Fig. 18b).

The genes involved in lignin synthesis had lower abundance than that in cellulose synthesis (Fig. 4a-b and Supplementary Table 15, 16). This indicated that the lignin synthesis ability was low, which was consistent with the low lignin content of the plant. Although some lignin synthesis related genes such as *PAL*, *4CL*, *F5H*, *COMT*, *CSE*, and *CCR* were expanded, their expression may be suppressed due to the regulation of other genes or transcription factors. There existed only two *CAD* genes in the synthesis of lignin monomers, which was one of the reasons limiting the synthesis of lignin. The expression level of *PAL* was high, but the substrate produced by *PAL* also existed in the synthesis pathway of flavonoids. Q-PCR results of some genes were consistent with the result of transcriptome profiling (Supplementary Fig. 15b). These results helped explain the genetic mechanism that elephant grass had a high herbage yield.

## Discussion

### The Assembly of an Allotetraploid Genome of Elephant Grass

The elephant grass genome (A’A’BB) is estimated to be highly heterozygous and 2.1-2.3 Gb in size. We assembled the 2.07 Gb chromosome-scale genome by integrating Nanopore, BioNano optical map and Hi-C technologies, with contig N50 of 2.9 Mb, scaffold N50 of 8.47 Mb, and 96.96% coverage of the full genome. This is the first chromosome-scale assembled genome of a tetraploid forage grass^15,40^. By reference to the genome of its closely related species *P. glaucum* (AA, 2n=14), the sequences were assembled into A’ and B sub-genomes each of which contained 7 pairs of collinear chromosomes. The quality of this assembly was significantly better than the recently published genome assemblies of orchard grass (*Dactylis glomerata* L.)^32^, pearl millet^18^, ryegrass (*Lolium perenne* L.), *T. urartu* and barley ^41^. Our sequencing and assembly strategy integrated Nanopore ultra-long reads sequencing, BioNano for chromosome-scale scaffolding, and a modified Hi-C protocol, which may be readily applicable to other complex genome sequencing and assembly^26^. The high-quality elephant grass genome sequence and transcriptome profiling data that we have completed and collected constituted important genetic, genomic and transcriptomic resources, which can be exploited in future research to understand evolution and genetic basis of complex traits such as biomass conversion and be used in molecular breeding.

### The Evolution of Species in *Pennisetum*

Based on the precision assembly of A’ and B sub-genomes of elephant grass, we reconstructed the phylogenetic tree of species in the grass family with a set of single-copy genes collinear in related species. The result showed that the species in *Pennisetum* diverged from *S. italica* approximate 22 MYA, and pearl millet (*P. glaucum*) and elephant grass (*P. purpureum*) were separated about 20 MYA. The allotetraploid elephant grass formed in the Miocene era of Africa about 15 MYA. Herbivorous pressure from herbivores mammals, particularly the Napieridae and Bovidae families, may have provided the impetus for the initial spread of C4 grasses across Africa continent during the Miocene era^42^. We inferred that the ancestorial A genome split into the ancestors of the elephant grass A’ and B genomes after *P. glaucum* differentiation. Afterward, the natural hybridization between A’A’ and BB diploid genomes produced the allotetraploid species several million years later. This inference was in part supported by GISH results done by ourselves (Supplementary Fig. 19) and the report on pearl millet and elephant grass^43^.

We also found the homologous relationship between chromosomes of related species and reasoned that the evolution of ancient chromosomes in the grass family was droved by double WGD events and chromosome fusion and lost events.Therefore, the high-quality genome of elephant grass is an enabling tool for understanding the evolutionary processes of *Poaceae* species and provide important genetic, genomic and transcriptomic resources for future community development.

### Genome Duplication and Significant Gene Expansion Laid the Foundation for the Giant Biomass of Elephant Grass

Our data and previous results showed that there were double ancient whole-genome duplication (WGD) events and polyploidization between A’ and B sub-genomes in elephant grass. These duplications and evolutionary selection pressure droved considerably the gene expansion and contract in the genome^44^. These may form the genetic foundation of its remarkable traits as fast growth, tolerance to abiotic stresses and high biomass accumulation^36,37^. Compared with pearl millet, there were not only thousands of gene families enhanced but also gene copy numbers expanded more in common gene families in A’ and B sub-genomes of tetraploid elephant grass. For example, the major gene families involved in cellulose and lignin biosynthesis pathways were significantly expanded with increased gene members. This was also confirmed by their highly abundant expression patterns in vegetative organs which were detected by RNA-Seq. This trend of gene family expansion following whole-genome duplication and polyploidization was also found in allohexaploidy common wheat, strawberry, and cotton^45^.

Furthermore, it is noticeable that functional differentiation existed between A’ and B sub-genomes. A’ sub-genome preferentially contributed to growth and development, whereas B sub-genome was mainly responsible for external stimuli and transportation. The high-quality reference elephant grass genome sequence reported here offered unprecedented insights into the genome evolution, polyploidization and genetic structure in these C4 fast growth grasses and paved a way for future studies, not only in elephant grass but also in other *Poaceae* species.

## Methods

### Plant Materials and Genomic in situ Hybridization (GISH)

The *Pennisetum purpureum* (access CIAT6263, Supplementary Fig. 1a) and *Pennisetum glaucum* (a hybrid) plant for GISH were collected from Jinan, Shandong Province, China. Genomic in situ hybridization had four steps: (1) Roots reproduced from stem cuttings of *P. purpureum* and seeds of *P.glaucum* were collected and treated; (2) Fixed root tips were digested for slides preparation; (3) Genomic DNAs of *P. glaucum* was labeled with biotin-16-dUTP by nick-translation reaction; (4) The hybridization for *P. purpureum* (as experimental) and *P. glaucum* (as control) were carried out according to the previous protocol^46^. Additional details are available in the Supplementary Note.

### Library Construction and Sequencing

Total genomic DNA was isolated from fresh leaves of elephant grass using QIAGEN® Genomic kit and purified from the gel by QIAquick Gel Extraction kit (QIAGEN). A total of 20 ug of high molecular weight DNA was used for Oxford Nanopore library and ultra-long reads library preparation (Supplementary Note). The purified library was loaded onto flow cells for sequencing on PromethION (Oxford Nanopore Technologies). Base-calling analysis of unprocessed data was performed using the Oxford Nanopore Albacore software (v2.1.3). After data quality control, 8.85 million Nanopore reads (~186.35 Gb data, ~93× coverage) and 1.70 million ultra-long reads (63.99Gb data, 32× coverage) were generated using Nanopore Technologies system. Nanopore reads had a mean length of 21.06 kb and N50 length of 28.48 kb, while ultra-long reads had a mean length of 37.63 kb and N50 length of 54.87 kb (Supplementary Table 1).

The high molecular weight DNA was extracted from fresh tissue for BioNano mapping. DNA labeling and staining were performed according to a protocol developed specifically by BioNano Genomics. Finally, 329.53 Gb of clean data were collected from BioNano Saphyr (Molecule >150 Kb; Molecule SNR > 2.75 & label SNR >2.75; Label intensity > 0.8). The average label number was 11.84 per 100 kb with an N50 size of 315.7 kb (Supplementary Table 1,3). For Hi-C sequencing, two libraries were prepared and sequenced on Illumina Novaseq 6000 to generate 1373.62 million (200.38Gb, 100× coverage) clean paired-end reads (Supplementary Table 1,3). Additional details are available in the Supplementary Note.

Total RNA was extracted from roots, stems, and leaves of all samples using the HiPure Plant RNA Kit according to the manufacturer’s instructions (Magen, Guangzhou, China). The PacBio Sequel platform (Pacific Biosciences, Menlo Park, CA, USA) was used for full-length RNA sequencing. A total of 1 mg of high-quality RNA from mixed tissues was used for library preparation with insert sizes of 0.5–4 kb and >4 kb (Supplementary Table 1). RNA-seq analysis was conducted using the Sequel platform according to the standard protocols (Supplementary Note). A total of 3 ug of RNA per sample was used for library preparation with insert sizes of 350bp and sequenced on Illumina Novaseq 6000. The full-length RNA-seq reads and RNA-seq reads of elephant grass were obtained for gene prediction analysis.

### Genome Assembly and Quality Assessment

The genome size was estimated before genome assembly. k-mer (k=17) analysis^47^ was performed using 2275.29 million 100bp paired-end reads. The genome size was thus estimated to be 2003.7 Mb (Supplementary Table 1, 2; Supplementary Fig. 2a). The DNA of elephant grass was also quantitatively analyzed by flow cytometry on MoFlo XDP (Beckman Coulter)^48^ and tomato^49^ was used as an internal reference, which suggested a genome size approximately 2.13 Gb for the haplotype (Supplementary Fig. 2b). Additional details are available in the Supplementary Note.

Nanopore reads and ultra-long reads were corrected using Nextdenovo (https://github.com/Nextomics/NextDenovo) and then used as the input for Smartdenovo (https://github.com/ruanjue/smartdenovo) assembly. The parameters for reads correction and assembly were as follows: read cuoff=1k, seed_cutoff = 13k, blocksize =1g; −k 21 −J 3000 −t 20. This resulted in the first assembly with a total size of ~2.22 Gb with Contig N50 of ~2.65 Mb (Table 1). After finishing the initial assembly, iterative polishing was conducted using Pilon (v1.22)^50^. The Pilon program was run with default parameters to fix bases, fill gaps and correct local misassemblies. Subsequently, the corrected genome (the size is ~2.27 Gb with Contig N50 of ~2.70 Mb, Table 1) was redundant using Redundans^51^ (https://github.com/lpryszcz/redundans) with the parameters as follows:--identity 0.9 --overlap 0.75. The final genome size is ~2.07 Gb with Contig N50 of ~2.90 Mb (Table 1). Finally, we performed BUSCO^29^ (embryophyta data set) assessments on the assembly. About 97.8% of the complete gene elements are found in the genome (Supplementary Table 4).

### Super Scaffold and Pseudomolecules Construction and Validation

In order to improve the quality of the genome assembly, single-molecule maps were de novo assembled into consensus maps using IrysView v2 software package (BioNano Genomics, CA, USA). The de novo genome assembly consisted of ~3,428.10 Mb of consensus genome maps (CMAP) with an average length of ~10.75 Mb and an N50 of ~44.02 Mb (Supplementary Table 1, 3). The assembly genome was compared with the optical genome map to correct anomalies. The genome scaffold was extended using Irys scaffolding with the default parameters. Overall, the elephant grass genome was assembled with ~2.08 Gb of scaffold sequences, had a larger scaffold N50 value of ~8.47 Mb while the longest scaffold was ~41.55 Mb (Table 1).

Hi-C technology is an efficient strategy for sequences cluster, ordered and orientation of pseudomolecule construction. Based on Hi-C data, ~175.69 million valid paired-end reads were used to assist genome assembly (Supplementary Table 1, 3). The genome sequence scaffolds and contigs were divided into subgroups, sorted and oriented into pseudomolecules using LACHESIS^52^ with the following parameters: CLUSTER MIN RE SITES = 100, CLUSTER MAX LINK DENSITY = 2.5, CLUSTER NONINFORMATIVE RATIO = 1.4, ORDER MIN N RES IN TRUNK=60, ORDER MIN N RES IN SHREDS=60. In the end, ~2006.94 Mb contigs were anchored to 14 pseudomolecules (Table 1).

The accuracy of the Hi-C assembly was evaluated by two methods. We first inspected the Hi-C contact heatmap. An elevated link frequency was observed with a diagonal pattern within individual pseudochromosomes, indicating the increased interaction contacts between adjacent regions (Supplementary Fig. 3). Additionally, the final assembly was mapped onto the pearl millet genome using NUCmer 3.1 (MUMmer v3.9.4alpha), the completeness comparison was performed by MUMmerplot 3.5 (MUMmer 3.23 package) on the NUCmer results after filtering to 1-on-1 alignments and allowing rearrangements with a 20 Kb length cutoff (Supplementary Fig. 4). The chromosomes were numbered according to the result of the mapped pearl millet genome. Additional details are available in the Supplementary Note.

### Genome Annotation

De novo repetitive sequences in elephant grass genome were identified using RepeatModeler (v1.0.4) (https://github.com/rmhubley/RepeatModeler) based on a self-blast search. RepeatMasker (v4.0.5) (http://www.repeatmasker.org/) was further adopted to search for known repetitive sequences using a cross-match program with a Repbase-derived RepeatMasker library, and the de novo repetitive sequences were constructed by RepeatModeler. The repeat-masked genome was used as input to two categories of gene predictors.

Protein-coding genes were predicted using a pipeline that integrated de novo gene prediction and RNA-seq-based gene models. For de novo gene prediction, Augustus (v3.0.3)^30^ and SNAP (v2006-07-28 https://github.com/KorfLab/SNAP) were run with default parameters and the training sets used were monocots and maize, respectively. For the RNA-seq-based prediction, 24 Gb RNA-seq reads from 3 tissues (root, stem, and leaf) and 10 Gb full-length RNA-seq reads were filtered to remove adaptors and trimmed to remove low-quality bases. Processed RNA-seq reads were aligned to the reference genome using TopHat2 (version 2.0.7)^53^. The transcripts were then assembled using Cufflinks (version 2.2.1)^54^

The rRNAs were predicted using RNAmmer (v1.2) ^55^, tRNAs were predicted using tRNAscan-SE (v1.23)^56^, and other ncRNA sequences were identified using the Perl program Rfam_scan.pl (v1.0.4) by inner calling using Infernal (v1.1.1)^57^.

Functional annotation of the protein-coding genes was carried out by performing BlastP (e-value cut-off 1e-05) searches against entries in both the NCBI and SwissProt databases. Searches for gene motifs and domains were performed using InterProScan (v5.28)^58^. The GO terms for genes were obtained from the corresponding InterPro or Pfam entry. Pathway reconstruction was performed using KOBAS (v2.0)^59^ and the KEGG database.

### Transcription Factor and Protein Kinases Annotation

iTAK program was applied to detect known TFs and PKs in elephant grass genome and other plant species^60^. The predicted gene sets were then used as queries in searches against the database. Finally, a total of putative TFs, belonging to 98 families and representing the predicted protein-coding genes, were identified; a total of 118 PKs families were identified.

### Transcriptome Sequencing

The stem (Supplementary Fig. 1c) segments of elephant grass growing for 40 days, 80 days and 120 days were collected and took the samples every other node from the root to the tip. In addition, young leaves, mature leaves (Supplementary Fig. 1b) and root (Supplementary Fig. 1d) of 120 days old elephant grass were collected and the leaves were divided into 3 parts: tip, middle, and base. All 19 samples were immediately frozen in liquid nitrogen after harvesting. Each sample had three biological replicates. RNA isolation, library construction, and sequencing were the same as RNA-seq, which were used for gene prediction analysis. Raw reads were trimmed to remove adaptors and enhance quality. Reads that were <100 bp in size after trimming were discarded. Overall, 106,967,926 (21.09 Gb, three replicate) to 162,667,319 (48.8 Gb, three replicate) raw reads were obtained for each sample (Supplementary Table 8).

The TopHat2 package (version 2.0.7)^53^ was used to map clean reads to the genome with the default parameters. Transcripts were assembled using Cufflinks (version 2.2.1)^54^. Gene expression was measured as transcripts per million reads (TPM) using Cufflinks. Differentially expressed genes (DEGs) were determined using DEseq^61^. The false discovery rate was used to adjust the P values. Genes with significant differences in expression, fold change > 2 and adjusted P-value <0.05, were considered as DEGs, and annotated to GO terms and KEGG pathways. Some of the genes related to the synthesis of cellulose and lignin were selected for expression verification by Q-PCR.

### Weighted Gene Co-Expression Network Analysis

Gene expression patterns for all identified genes were used to construct a co-expression network using WGCNA (v.1.47)^39^. Genes without expression detected in all tissues were removed before analyses. Soft thresholds were set based on the scale-free topology criterion^62^. An adjacency matrix was developed using squared Euclidean distance values, the topological overlap matrix was calculated for unsigned network detection using the Pearson method. Co-expression coefficients >0.55 between the target genes were then selected. Finally, the network connections were visualized using Cytoscape^63^.

### Expression Bias of Homologous

Protein-coding genes from A’ and B sub-genomes of elephant grass were employed as queries in a BLAST search against each other. The best reciprocal hits with > 80% of identity, an E-value cutoff of <1E-30, and an alignment accounting for >80% of the shorter sequence were obtained as gene pairs between A’ and B sub-genomes. On the best reciprocal BLAST matches between A’ and B sub-genomes, we identified 11,257 gene pairs that had a 1:1 correspondence across the two homologous sub-genomes. To investigate the expression bias of these paired homologous from the two sub-genomes, we calculated the TPM values of the homologous in root, stem, and leaf. DEGs were determined using DEseq^61^.

### Syntenic and Ks Analysis

Syntenic blocks were identified using MCScanX with default parameters^64^. Proteins were used as queries in searches against the genomes of other plant species to find the best matching pairs. Each aligned block represented an orthologous pair derived from the common ancestor. Ks (the number of synonymous substitutions per synonymous site) values of the homologous within collinear blocks were calculated using Nei-Gojobori approach implemented in PAML^65^, and the median of Ks values was considered to be the representative of the collinear blocks. The values of all gene pairs were plotted to identify putative whole-genome duplication events within elephant grass. The duplication time was estimated using the formula t = Ks/2r, where r is the neutral substitution rate, to estimate the divergence time between two sub-genomes and other species. A neutral substitution rate of 8.12 × 10^−9^ was used in the current study.

### Phylogenetic Tree Construction and Evolution Rate Estimation

Orthologous gene clusters in A’ and B sub-genomes of *P. purpureum* and 11 other representative plants (Supplementary Table 10) were identified using OrthoMCL program^34^. A total of 15,546 homologous groups containing 6,926 genes specific to elephant grass were identified with a total of 169 single-copy orthologous in this set. The single-copy orthologous genes were used to build an ML tree by FastTree (v2.1.9)^66^. This ML tree was converted to an ultrametric time-scaled phylogenetic tree by r8s (Sanderson, 2003) using the calibrated times from the TimeTree^67^ website. For example, 120.0–155.8 Mya for *A. thaliana* and rice, 39.4–53.8 Mya for rice and *B. distachyon* and 22.7–28.5 Mya for *S. italica* and *S. bicolor*.

### Identification of Chromosome Reshuffling

After identification of the syntenic and colinear blocks between elephant grass and other grasses, we set the rice genome as the reference and identified all the homologous chromosome relationships between rice and A’ and B sub-genomes of elephant grass, rice and *P. glaucum*.

### Gene Family Analysis

Gene family expansion or contraction was determined using CAFE’(v3.0) ^68^. The gene families that were identified in at least 5 species were selected for further analysis. A random birth-and-death model was used to evaluate changes in gene families along each lineage of the phylogenetic tree. A probabilistic graphical model (PGM) was used to calculate the probability of transitions in each gene family from parent to child nodes in the phylogeny. The extensional and contractile family of all nodes and species were analyzed.

To investigate the genes involved in the cellulose and lignin biosynthesis pathways in *P. purpureum* genome, genes were detected by the annotation result. We retrieved protein sequences of these gene families from elephant grass for homology-based searches with the criteria of similarity >80% and coverage >80%. We then confirmed the presence of the conserved domain within all protein sequences and removed members without a complete domain. Protein domains of these homologs were predicted by Pfam (http://pfam.xfam.org/). Only the genes with the same protein domain were considered as homologs.

## Data Availability

The elephant grass genome has been deposited under BioProject accession number PRJNA607017.

## Acknowledgments

The Nanopore, BioNano and Hi-C sequencing and primary assembly were performed with the help of the Nextomics Biosciences Institute in Wuhan, China. This study was financially supported by the Integration of Science and Education Program Foundation for the Talents by Qilu University of Technology (No. 2018-81110268), Foundation of State Key Laboratory of Biobased Material and Green Papermaking (No. 2419010205 and No. 23190444).

## Author Contributions

T.X., W.W. designed the project and contributed the original concept of the manuscript. S.Zhang and Z. Xia performed *de novo* genome assembly and annotation, analyzed the data as a whole and wrote the manuscript. W. Zhang and X. Zhao completed the Q-PCR validation of selected genes. C. Li, X. Wang, X. Lu, H. Ma performed DNA, RNA extraction and cytogenetics study. W. Zhang participated in the interpretation of the data and revised the manuscript. X Zhou conducted the repetitive sequence analysis. T. Zhu assisted in BioNano research. G. Liu, P Liu and H.Yang performed the biological characteristics study. J. Arango and M. Peters provided data and information of growth and planting of elephant grass and reviewed the manuscript.

## Competing Interests

The authors declare no competing interests.

